# Internal oligo(dT) priming in bulk and single cell RNA sequencing

**DOI:** 10.1101/2021.09.24.461289

**Authors:** Marek Svoboda, H. Robert Frost, Giovanni Bosco

## Abstract

Significant advances in RNA sequencing have been recently made possible by the use of oligo(dT) primers for simultaneous mRNA enrichment and reverse transcription priming. The associated increase in efficiency has enabled more economical bulk RNA sequencing methods as well as the advent of high throughput single cell RNA sequencing, now already one of the most widely adopted new methods in the study of transcriptomics. However, the effects of off-target oligo(dT) priming on gene expression quantification have not been fully appreciated. In the present study, we describe the extent, the possible causes, and the consequences of internal oligo(dT) priming across multiple publicly available datasets obtained from a variety of bulk and single cell RNA sequencing platforms. In order to explore and address this issue, we developed a computational algorithm for identification of sequencing read alignments that likely resulted from internal oligo(dT) priming and their subsequent removal from the data. Directly comparing filtered datasets to those obtained by an alternative method reveals significant improvements in gene expression measurement. Finally, we infer a list of genes whose expression quantification is most likely to be affected by internal oligo(dT) priming.

## Introduction

Next generation sequencing (NGS) of RNA is one of the most commonly used methods to measure gene expression in tissues as well as single cells, yielding readily quantifiable information about the relative levels of protein production as a proxy to cellular activity. However, protein-coding messenger RNA (mRNA) constitutes only a relatively small portion of total RNA, the majority being ribosmal RNA (rRNA) by mass and transfer RNA (tRNA) by number of molecules (1). The RNA subset of interest is therefore usually obtained by either rRNA depletion (negative selection via rRNA-specific probe hybridization) or mRNA enrichment (positive selection via oligo(dT) hybridization to mRNA poly(A) tails; 2). The latter is frequently preferred in studies focusing on proteincoding genes, where specificity for mRNA is desired for higher processing efficiency and reduced sequencing cost (3).

RNA sequencing library preparation then proceeds with cDNA synthesis by supplying reverse transcriptase and short DNA oligo primers annealed to the RNA of interest. This can be achieved through random oligo priming, resulting in relatively unbiased sequencing reads from across the entire length of mRNA (4, 5). Alternatively, oligo(dT) probes may be used directly to prime cDNA synthesis, while also enriching for poly(A) tail containing mRNA in the same step. While the oligo(dT) method tends to yield sequencing reads biased towards the transcripts’ 3’ end, its simultaneous mRNA enrichment and cDNA priming is especially useful for gene expression quantification using very low abundance input RNA, for example from individual cells (6). For this reason, oligo (dT) priming is employed by virtually all existing high-throughput single cell RNA sequencing (scRNA-seq) methods (e.g. Drop-seq, inDrop, 10X, Seq-well, sci-RNA-seq; 7–11), some of the low-throughput ones (e.g. Smart-seq, CEL-seq; 12, 13), and even some costeffective bulk RNA sequencing methods (e.g. QuantSeq, TagSeq, 3’Pool-seq; 14–16). Most of these methods aim to quantify gene expression by counting the number of mRNA molecules that originate from their respective genes using the underlying assumption that oligo(dT) priming only takes place at the 3’-poly(A) tails. The number of priming events is therefore assumed to correspond to the number of original molecules of mRNA before any subsequent PCR amplification takes place. However, this assumption is only correct for transcripts that do not contain additional, internal oligo(dT) priming sites.

While the 3’-terminal poly(A) tails of mRNA are added during post-transcriptional processing of nascent mRNAs, some of the RNA molecules also contain genome-encoded poly(A) sequences (adenine single-nucleotide repeats or A-SNRs). These internal priming sites can lead to “off-target” oligo(dT) hybridization, as reported previously in studies on gene identification, alternative polyadenylation, and scRNA-seq analysis (Figure 1A). Early attempts at gene identification that relied on 3’-terminal priming of entire transcripts often resulted in truncated cDNAs whereby internally hybridized oligo(dT)s initiated cDNA synthesis and simultaneously prevented extension of reverse transcription initiated at a downstream poly(A) tail (17). Through this process, internal priming led to misidentification of the extent of transcripts. Additionally, it has been suggested that internal oligo(dT) priming (onto genome-encoded A-SNRs) may take place more efficiently than terminal priming (onto poly(A) tails). The recommended solution is the use of anchored oligo(dT) primers (i.e., oligo(dT)s with 1-2 nucleotides other than T at their 3’ end), which limits but does not completely prevent internal oligo(dT) priming. Similarly, in studies that focus on alternative polyadenylation, internal oligo(dT) priming may lead to identification of false polyadenylation sites (18, 19). In this context, various rules and methods have been previously used in downstream data analysis to identify and filter out sequencing read alignments that might have resulted from internal oligo(dT) priming. Examples include identification of the following genomic features adjacent to the 3’ end of the alignments, suggestive that the sequencing reads originated from internal oligo(dT) priming: six continuous adenines (As) or more than seven As in a 10 nt window (18), AG-runs of six or more nucleotides or eight or more As or Gs in a 10 nucleotide window (20), eight or more As or high A/T content (27 out of 30 bases; 21), and 12 or more adenines present in an 18 nucleotide window (22). Although these criteria have been used repeatedly in the alternative polyadenylation literature to identify internally primed alignments, they are seemingly arbitrary and their efficacy has not been ostensibly demonstrated. In the context of scRNA-seq, internal oligo(dT) priming has only been previously discussed for the purposes of an analysis termed “RNA velocity” (23). Because A-SNRs are commonly found in introns, oligo(dT) hybridization to these sequences in unspliced pre-mRNA may lead to generation of intronic sequencing reads. Therefore, in RNA velocity, detection of internally primed intronic alignments in scRNA-seq data is used to derive changes in gene expression over time. Additionally, a bias in sequencing data associated with oligo(dT)s hybridizing to genomic poly(A) sequences has been previously reported in the SMART library preparation method (24), which is often used in scRNA-seq to yield complete coverage across transcripts, but the impact on the measured gene expression remains unclear.

**Figure 1.**
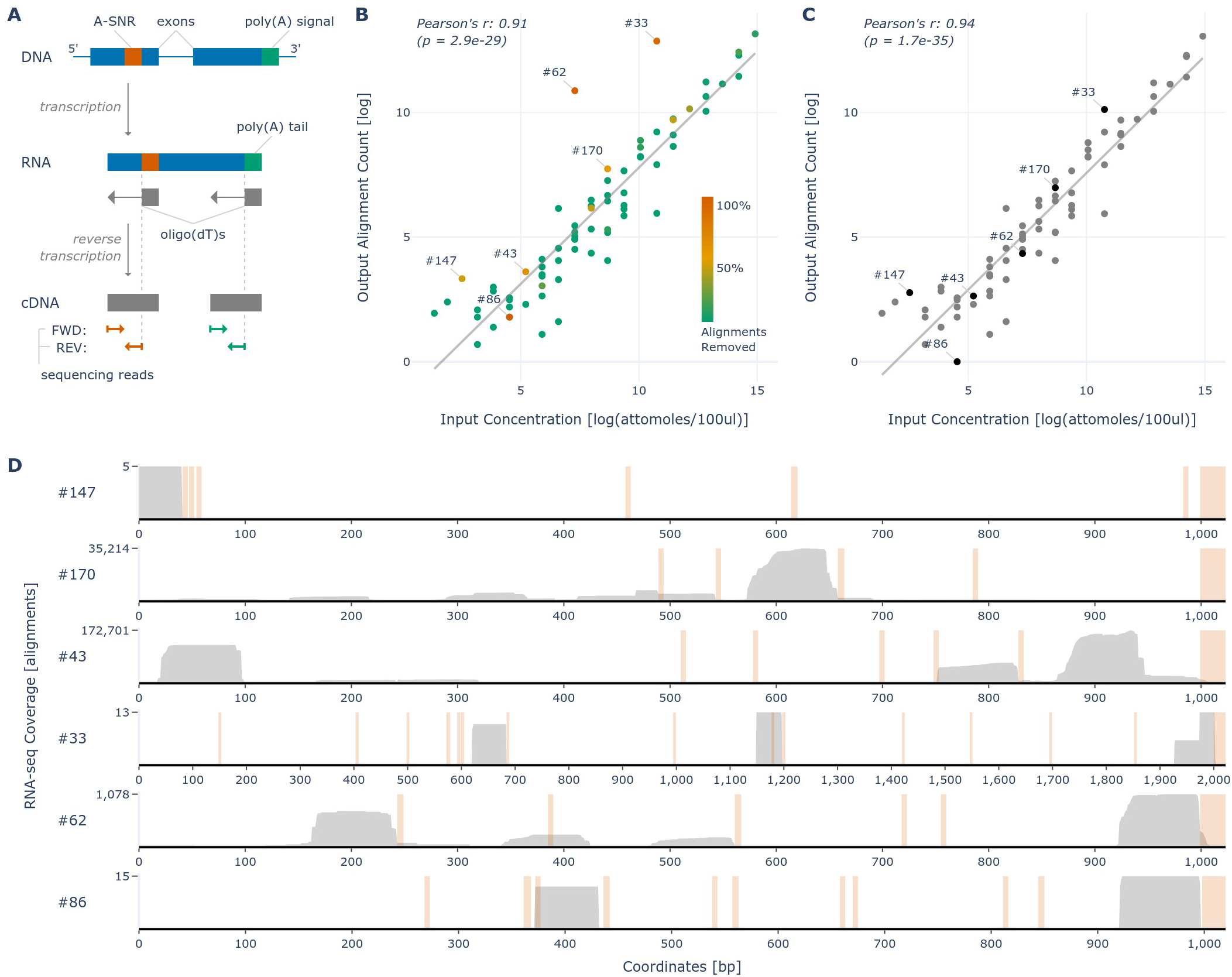
Removal of non-terminal ERCC alignments from bulk oligo(dT) priming-based data. **A.** A simplified diagram describing the overall steps in oligo(dT)-primed RNA sequencing library preparation and sequencing process. Depending on the sequencing method used, the sequencing takes place either from the 5’ (FWD methods used for mRNA quantification) or the 3’ (REV methods used for polyadenylation site detection) end of the cDNA fragment, resulting in reads in either sense or anti-sense direction, respectively. For that reason, each sequencing read in the REV methods is directly adjacent to the oligo(dT) priming event that gave rise to its cDNA, while in the FWD methods, each sequencing read maps up to about 500 bp (given by the length of the cDNA fragments sequenced) upstream of its priming event. **B.** Correlation between log of input ERCC concentrations and log of the respective alignment counts detected using oligo(dT)-primed bulk RNA sequencing; data by Wu, Schmid, Rib et al. (26). ERCCs not detected are not shown. The color scale indicates the proportions of non-terminal reads (more than 75 nt upstream of the poly(A) tail) aligned to each respective ERCC reference. Numbers of the six ERCCs with the highest proportions of non-terminal coverage are annotated. **C.** Correlation plot similar to that in **B.** after the removal of non-terminal alignments, with the same ERCCs annotated. **D.** The RNA sequencing coverage tracks of the six ERCCs annotated in **B.** and **C.** Per-basepair sequencing coverage is shown in gray with the maximum coverage indicated on the y-axis. In orange, SNRs of five or more consecutive As (including the poly(A) tails, which are synthesized as a part of the ERCCs) in each ERCC reference are highlighted.

To our knowledge, internal oligo(dT) priming to A-SNRs has not been thoroughly characterized nor addressed in the context of gene expression quantification using oligo(dT)-based RNA sequencing. The fact that oligo(dT) primers may spuriously hybridize to one or more internal, genome encoded A-SNRs in addition to the targeted poly(A) tails of mRNA molecules from a given gene could lead to a higher probability or even multiplicity of detection of mRNA molecules that contain these A-SNRs. We therefore reasoned that internal oligo(dT) priming may lead to relative overinflation of measured mRNA molecule abundance in genes that contain genome-encoded poly(A) sequences, introducing bias in the gene expression data.

We analyzed previously existing publicly available datasets obtained with various bulk and single cell RNA sequencing methods that use oligo(dT) priming to count mRNA molecules and found that all of them exhibited signs of internal priming. We subsequently developed an algorithm to detect genome-encoded A-SNRs as well as the sequencing read alignments that likely resulted from internal oligo(dT) priming at these sites in mRNA. We show that removing these internally primed alignments as an additional data processing step improves the accuracy of gene expression count data. Based on these findings, we ranked human genes by their likelihood to be subject to internal oligo(dT) priming when expressed.

## Results

### External RNA Controls Consortium (ERCC) spike-in standards exhibit signs of internal oligo(dT) priming

ERCC standards are synthetic RNA molecules with poly(A) tails that are routinely used for baseline measurements and normalization of RNA sequencing data by comparing the known input ERCC concentrations to the sequenced output quantities (25). In order to assess whether oligo(dT)s may be prone to internal priming, we first set out to examine a publicly available dataset by Wu, Schmid, Rib et al. (26), which was enriched with ERCC spike-in standards. This dataset was generated using the QuantSeq 3’ mRNA-Seq REV library prep kit, which is a bulk RNA sequencing protocol that aims to pinpoint the exact 3’ ends of transcripts by generating antisense reads directly upstream of the oligo(dT) priming event (Figure 1A). We reasoned that if internal priming did occur during the creation of this dataset, we would be able to detect outliers among the ERCC standards whose measured output alignment counts are higher than expected from their input concentrations.

Indeed, by comparing the known input concentrations of the ERCC standards to their respective resulting alignment counts after sequencing, we detected several notable outliers in the data (Figure 1B). After removal of all non-terminal ERCC alignments, which mapped further (i.e. more than 75 basepairs (bps)) upstream than reasonably expected to result from the oligo(dT)s priming onto the poly(A) tails, the outliers’ alignment counts moved closer to their expected values and the overall correlation of the input concentrations with the detected output counts increased significantly (Pearson’s r increased from 0.91 to 0.94, *p* ≈ 0.006; Figure 1C). We examined where the alignments mapped on the ERCC reference sequences and found that a large portion of the ERCC-associated non-terminal alignments mapped directly upstream of internal SNRs of five or more consecutive As (Figure 1D). ERCC spike-in standards therefore may be subject to internal oligo(dT) priming, possibly secondary to oligo(dT)s’ hybridization to A-rich internal sequences on the ERCC standards. Interestingly, Wu, Schmid, Rib et al. did not use the ERCC sequencing data in their final data analyses, citing inconsistent results.

### Oligo(dT)-primed RNA sequencing reads are enriched upstream of genome encoded poly(A) sequences

To determine if internal priming might also be detected in experimental RNA sequencing data, we first scanned the human genome for the presence of A-SNRs and quantified their relative abundance by length (Figure 2A; for scanning results of other commonly used reference genomes, see Figure S1), as reported previously (27). Interestingly, the relative frequency of A-SNRs nine or more nucleotides long normalized by their expected frequency increased logarithmically with increasing A-SNR length, suggesting the presence of evolutionary processes that favor long stretches of adenines. Since most RNA sequencing data analyses commonly focus on reads that specifically align to exonic sequences of genes, we also determined the relative distribution of A-SNRs with respect to their genomic annotations (Figure 2B and S1). We found that while the abundance of A-SNRs was relatively depleted in exons compared to other genomic annotations, exons still contained a considerable proportion of A-SNRs, rendering these sites viable candidates for internal oligo(dT) priming that could potentially result in inflated exonic sequencing read counts.

**Figure 2.**
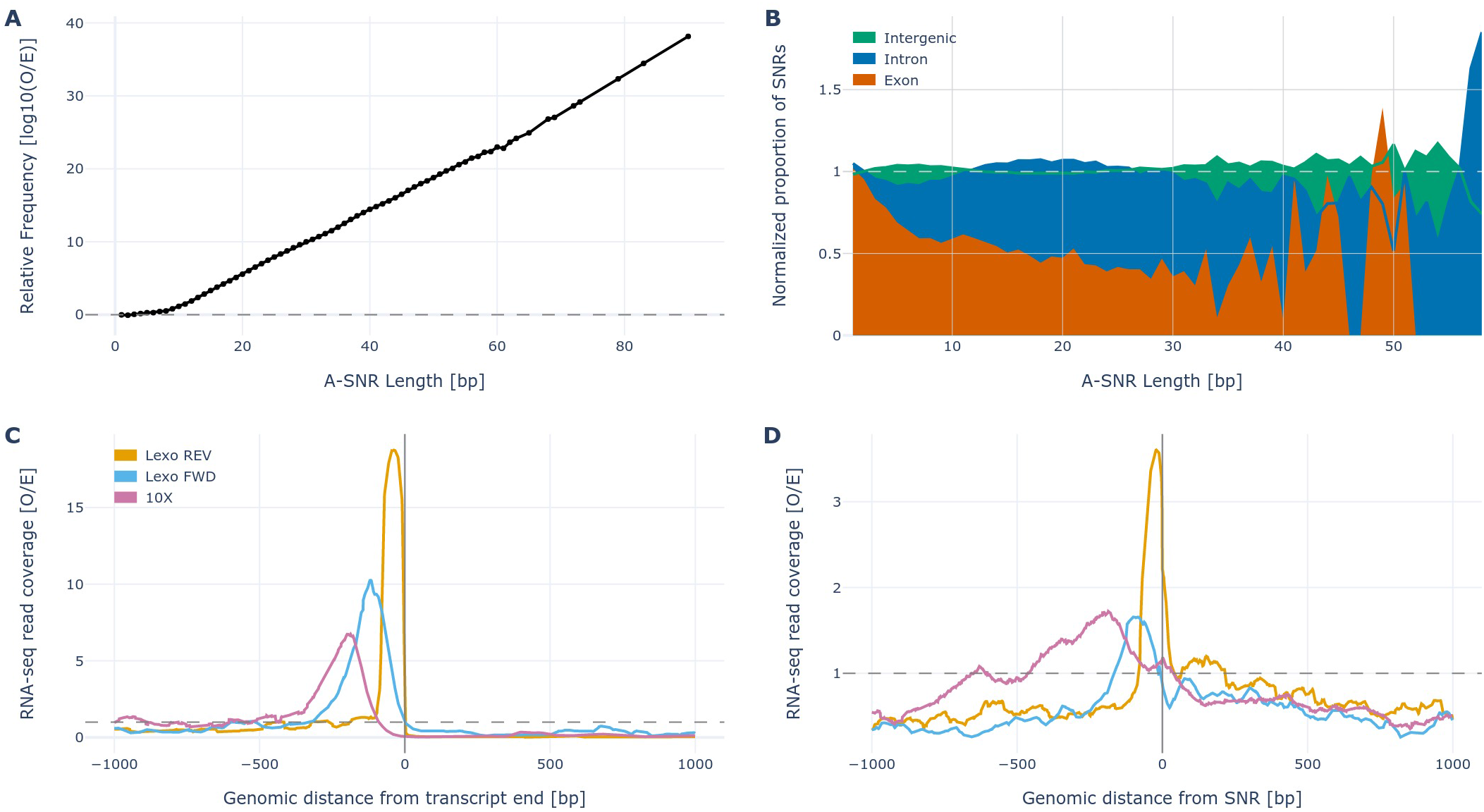
Genome encoded A-SNRs and the associated sequencing coverage. **A.** Log10 of relative (observed/expected) frequency of A-SNRs by length across the entire human genome reference. In gray dashed line, the expected frequency represents the probability that A-SNRs of given length be found by pure chance. **B.** The relative enrichment of A-SNRs of each length by their genomic regions, normalized by the respective proportions of the human genome thus annotated. In gray dashed line, the expected normalized proportion of A-SNRs for each region represents random allocation of A-SNRs across the genome. SNR lengths represented by fewer than 10 SNRs (i.e. longer than 58 bp) were truncated. **C.** Normalized aggregate stranded exonic RNA sequencing coverage in the vicinity of all annotated transcript 3’ ends, which are aligned at “0 bp” in the sense orientation. **D.** Normalized aggregate stranded exonic RNA sequencing coverage in the vicinity of all A-SNRs of five nucleotides or longer, whose starts (5’ ends) are aligned at “0 bp” in the sense orientation. In both **C.** and **D.**, sequencing coverage from datasets by Wu, Schmid, Rib et al. (“Lexo REV”; 26), Ma et al. (“Lexo FWD”; 5), and Ding et al. (“10X”; 28) is depicted and gray dashed line represents the expected coverage, i.e. aggregate exonic sequencing coverage in the dataset divided by the total length of all collapsed exons in the respective genome.

We therefore measured aggregate exonic RNA sequencing coverage upstream of potential priming events in three datasets, each created using a different library preparation method but all utilizing oligo(dT) priming: a QuantSeq 3’ mRNA-Seq REV dataset by Wu, Schmid, Rib et al. discussed above (“Lexo REV,” bulk RNA sequencing in the anti-sense orientation; 26), a QuantSeq 3’ mRNA-Seq FWD dataset by Ma et al. (“Lexo FWD,” bulk RNA sequencing in the sense orientation; 5), and a 10X Genomics Chromium Single Cell 3’ (v2) dataset by Ding et al. (“10X,” scRNAseq in the sense orientation; 28). First, we measured aggregate exonic RNA sequencing coverage aligned in the vicinity of annotated transcripts’ 3’ ends (terminal coverage) across the respective reference genomes. As expected, we observed an enrichment of sequencing reads directly upstream (Figure 2C) of transcript 3’ ends, originating from terminal oligo(dT) priming onto poly(A) tails of mRNA molecules. Also as expected from the differences between the library preparation methods (Figure 1A), alignment enrichment in the Lexo REV dataset was found directly upstream of and adjacent to the transcript 3’ ends (reads being in the anti-sense orientation with respect to the transcript reference), while the enrichment peaks were further upstream in the Lexo FWD and 10X datasets (with reads in the sense orientation). Analogously, we scanned the same datasets for aggregate exonic coverage in the vicinity of exonic SNRs of five or more continuous adenines (internal coverage) across the respective genomes and intriguingly, we also found enrichment directly upstream of these loci and the distances of the enrichment peaks from the A-SNRs mirrored those seen upstream of transcript 3’ ends, as described above (Figure 2D). The most likely explanation for this internal coverage enrichment is that these sequencing reads resulted from oligo(dT)s priming onto the A-SNRs found in the sequenced mRNA molecules. Moreover, the relative enrichment upstream of A-SNRs increased with increasing A-SNR length, suggesting that longer stretches of A-SNRs are more likely to attract oligo(dT) priming (Figure S2).

### Removal of internally primed reads improves the accuracy of oligo(dT)-based RNA sequencing data

We next sought to investigate whether removal of internal alignments upstream of A-SNRs may lead to improved accuracy of mRNA counts in oligo(dT)-primed RNA sequencing data. For this purpose, we focused on analysis of publicly available datasets that had been generated using the most common bulk and single cell RNA sequencing library preparation methods for counting of mRNA molecules via oligo(dT) priming and sense-oriented sequencing (QuantSeq FWD bulk dataset by Ma et al. (5), and 10X, CEL-seq2, Drop-Seq, inDrop, and Seq-well single cell datasets by Ding et al. (28)). Each of these datasets also had an associated random oligo-primed bulk RNA sequencing dataset generated from the same biological sample used as a gold standard. Due to the constraints of next generation sequencing, all of these methods are optimized to yield cDNA libraries with fragments of length limited to about 500 bp (e.g., the mean fragment size including adaptors for the QuantSeq libraries is 335-456 bp, with the cDNA insert size being 203-324 bp; 29). In oligo(dT) priming-based methods, the reads that align to the reference genome are sequenced from the 5’ end toward the poly(A) tail in the sense strand (FWD) orientation and are usually less than 100 bp long (Figure 1A). As a result, these sequencing reads align on the reference genome up to several hundred bps upstream of the original oligo(dT) priming event (either onto a poly(A) tail or an A-SNR), complicating the association of the priming event with the resulting sequencing read alignment. It is notable that this distance gap between the priming event and sequencing read alignment is observed in methods that focus on mRNA quantification; those that instead focus on polyadenylation site detection commonly use antisense-oriented (REV) sequencing reads directly upstream of and adjacent to the oligo(dT) priming event, like in the QuantSeq 3’ mRNA-Seq REV dataset by Wu, Schmid, Rib et al. (26) discussed above. Therefore, in order to identify and remove the sequencing read alignments that likely resulted from internal (A-SNRs) priming in RNA counting FWD library preparation methods, the correct distance of the alignments from their respective priming events (*maximal coverage distance*) has to be determined.

In addition to the *maximal coverage distance,* the minimal length of an A-SNR that is sufficient for oligo(dT) priming to occur (*minimal A-SNR length*) also has to be determined for correct identification of internally primed alignments. Although several different cutoffs have been used in the past for these purposes, none of them had been previously validated (18, 20–22). Finally, as we suspected that A-SNRs may attract oligo(dT) priming even if they contain one or several other bases, we also had to determine the *maximal number of mismatches* allowed in A-SNRs for internal priming in these library preparation methods.

We therefore developed an algorithm that filters an aligned RNA sequencing dataset given the parameters of *maximal coverage distance, minimal A-SNR length*, and the *maximal number of mismatches* allowed. By iteratively applying this filtering algorithm, we then optimized these three parameters separately for each oligo(dT) priming-based dataset studied to maximize its accuracy, i.e. its correlation with a gold standard bulk RNA sequencing dataset (one that uses random oligo priming) obtained from the same biological sample (Figure S3). We found that the optimal parameters as well as the relative increase in correlation differed across the methods but for each of the datasets studied, such combination of filtering parameters could be found that resulted in a statistically significant increase in RNA sequencing accuracy (Figure 3A). Although removal of all non-terminal alignments improved the data accuracy in all non-bulk datasets for at least some values of *maximal coverage distance*, it is important to note that in all the datasets studied the highest increase in accuracy was observed after specific removal of only those alignments that were associated with A-SNRs. Additionally, there was no correlation between the amount of exonic alignments removed and the resulting accuracy of the dataset (Figure S4). When a proportion of exonic alignments equal to that in the optimally filtered dataset was removed randomly (regardless of their association with A-SNRs), the accuracy of the dataset decreased compared to the baseline before filtering (Figure S4D). These results support the idea that internal priming events from A-SNRs specifically contribute to erroneous sequencing reads and thus decrease the accuracy of gene expression quantification.

**Figure 3.**
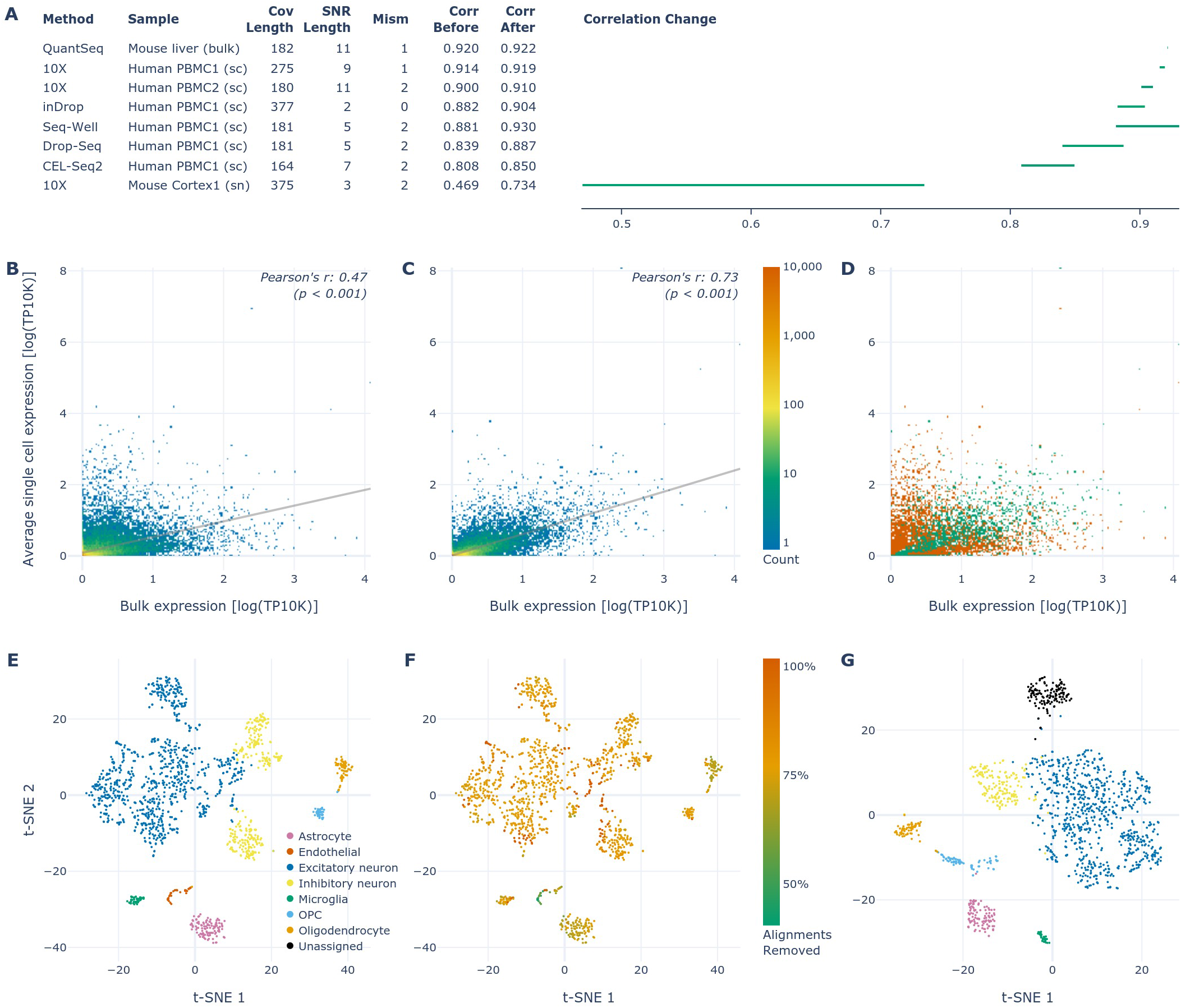
Optimized filtering parameters by library preparation method and details of the single nuclei dataset. **A.** Table of the optimized filtering values for each respective dataset studied along with the correlation change (visualized difference between the correlation before and after filtering; single cell correlation values before filtering reproduced from Supplementary Figure 11 in 28). *p* << 0.001 for each individual correlation, as well as for each difference between the correlations before and after filtering (adjusted using the Bonferroni correction). Abbreviations used: “Cov Length”: maximal coverage distance; “SNR Length”: minimal A-SNR length; “Mism”: maximal number of mismatches; “Corr”: correlation; “sc”: single cells; “sn”: single nuclei. **B.** Correlation of gene expression in the sample of mouse cortex nuclei between the 10X (oligo(dT)) and bulk (random oligos) methods, with intronic alignments included in gene expression quantification for both. **C.** Correlation between the same datasets as in **B.** after filtering the oligo(dT) dataset using the optimal parameters for this dataset listed in **A.** Both **B.** and **C.** are 2D histograms sharing the rainbow color scale depicting the density of genes in each region with the line of best fit in gray. **D.** Visualization of changes between **B.** and **C.**, where the regions with fewer and more genes in **C.** relative to **B.** are depicted by orange and green, respectively. **E.** t-SNE plot of the single nuclei mouse cortex 10X dataset colored by the cell types detected (reproduced from Figure 6A in 28). **F.** The same t-SNE plot as in **E.**, colored by the proportion of alignments optimally filtered out from each cell, as per the adjacent color scale. **G.** t-SNE plot of the same dataset as in **E.** and **F.**, after the internally primed alignments have been optimally filtered out. The colors correspond to the same cell types as in **E.**

The improvement of accuracy was the highest for the single nuclei dataset, where sequencing reads aligned to introns (along with those aligned to exons) were included in gene expression quantification (Figure 3B-G), as is common for single nuclei sequencing datasets (28, 30). As expected from the higher relative abundance of A-SNRs in introns than in exons (Figure 2B), this dataset had the highest proportion of alignments filtered out, removing about 64% of intragenic (exonic and intronic) alignments overall (Table S1), ranging from 38% to 100% per cell (Figure 3F). Interestingly, after filtering, the “Endothelial” cell type was no longer detected in this mouse cortex sample, while one of the two cell clusters previously assigned to the “Inhibitory neuron” cell type was now “Unassigned,” confirming the spatial separation of these two cell clusters in the t-SNE plot as indicative of a difference in cell types (Figure 3G). When we analyzed this dataset considering only exonic alignments (as is common for single cell datasets), the correlation with the associated bulk dataset was significantly higher than with intronic alignments included before filtering (0.710 without, compared to 0.469 with intronic alignments included) and it further increased after optimal filtering (0.753 compared to 0.734, respectively; Figure S5). Inclusion (or lack thereof) of intronic alignments in the analysis of each associated random oligo bulk dataset mirrored that of the oligo(dT) dataset it was compared to. Because introns contain a relatively high abundance of A-SNRs, which are associated with spurious sequencing reads, inclusion of intronic alignments likely further decreases the accuracy of gene expression quantification.

### Linear model ranks genes by the likelihood that their reads result from internal oligo(dT) priming

Of the methods studied, 10X is currently the most widely used mRNA counting library preparation method, as it is the most user-friendly and it shows most consistency in results among the single cell methods (28). Additionally, its accuracy was the highest at baseline before the filtering and also the *minimal A-SNR length* for optimal filtering was the highest among the single cell methods tested, resulting in the lowest proportion of alignments removed (Table S1). Therefore, we used this dataset to derive a conservative linear model to predict the proportion of sequencing reads optimally filtered out for each gene as a function of the exonic A-SNRs it contains. Using these predictions, we ranked all genes by the likelihood that their alignments result from internal oligo(dT) priming (Table S2).

The ten genes with the highest predicted probabilities of internal oligo(dT) priming all contain numerous A-SNRs of lenghts nine or higher with up to one mismatch (Figure 4). All of these genes were detected as expressed in the 10X single cell dataset, with only a minority of the sequencing reads aligned in the terminal portions of annotated transcripts (i.e. upstream of the transcripts’ 3’ ends), while most of the alignment peaks were found upstream of the genome-encoded A-SNRs mentioned above. This observation strongly suggests that these alignments originated from internal oligo(dT) priming. Although these data represent sequencing of single cell RNA (as opposed to that from single nuclei, which is expected to contain a higher proportion of intronic sequences) and the gene predictions were made with respect to only exonic A-SNRs, the top ten genes show a considerable amount of intronic coverage as well, in agreement with previous studies (23). Interestingly, cells expressing the highest ranked gene, KCNQ1OT1, have been previously discarded from oligo(dT) priming-based scRNA-seq datasets due to “artifactual” expression of this gene (31). Filtering out internally primed alignments may have obviated the need to discard cells from these scRNA-seq datasets.

**Figure 4.**
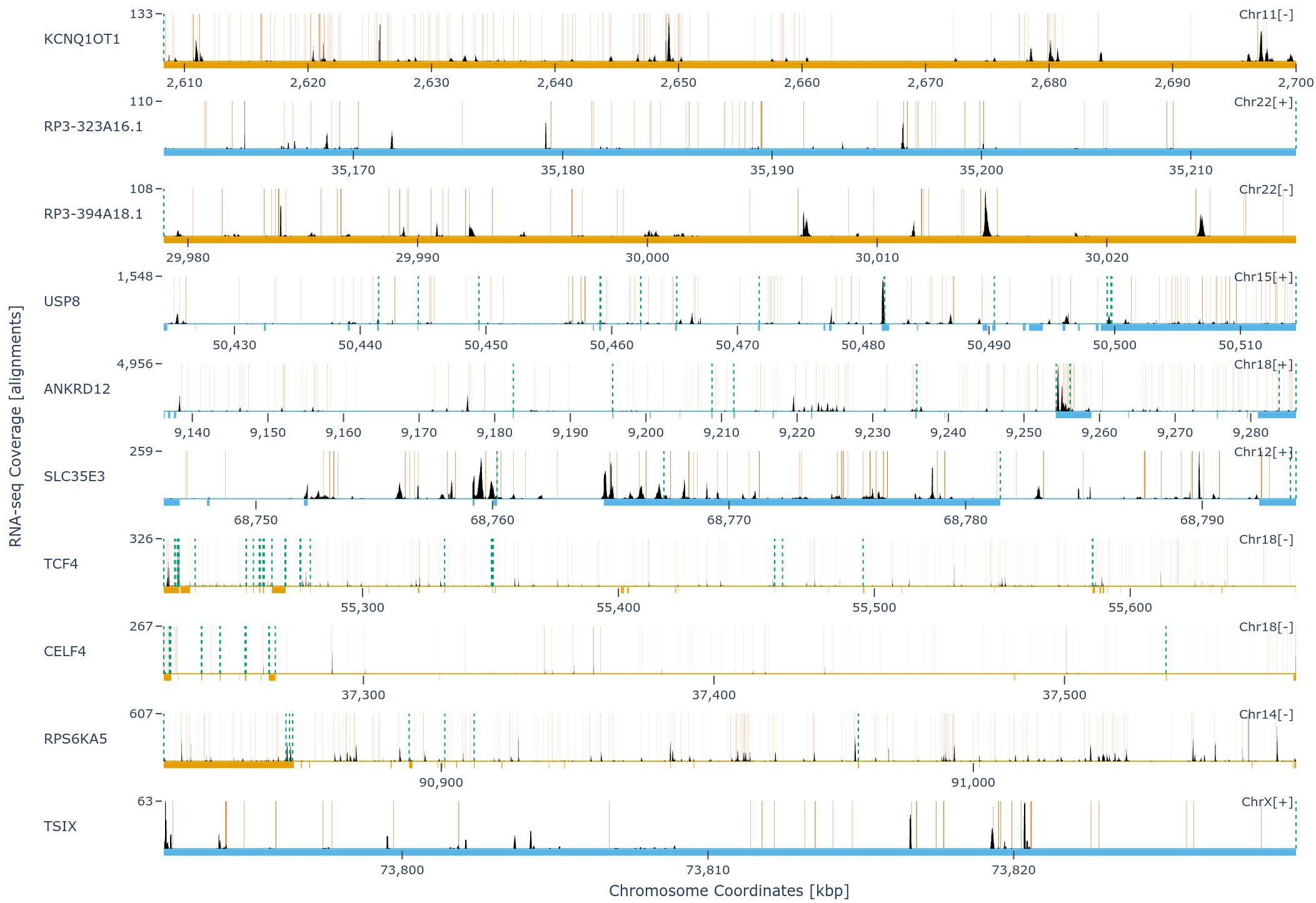
Sequencing coverage of the top 10 genes predicted to be affected by internal oligo(dT) priming. Genome tracks of each respective gene with the chromosome coordinates (x-axes) shown in the “+” strand orientation. The sense strand of each gene is indicated next to the chromosome number (“Chr#[+/-]”) in the upper right corner of each genome track and in color of the track with light blue signifying the “+” strand (displayed in 5→3’ orientation) and light orange color signifying the “-” strand (displayed in 3→5’ orientation). The thick bars underneath the genome tracks depict the extent of exon annotations collapsed across all transcripts for each respective gene. Per-basepair sequencing coverage is shown in black with the maximum coverage indicated on the y-axis. The locations of genome-encoded SNRs of nine or more adenines with up to one mismatch are indicated by the solid orange vertical lines. The different 3’ ends (polyadenylation sites) for each mRNA transcript variant are indicated by the dashed green vertical lines. In the absence of internal poly(dT) priming, sequencing coverage would be expected to be detected only upstream (in each gene’s sense orientation) of these 3’ ends.

## Discussion

Oligo(dT) priming-based sequencing has revolutionized biology in the recent years by enabling massively parallel sequencing of RNA from individual cells, while also inspiring the emergence of more economical bulk RNA sequencing methods. As next generation sequencing is being gradually replaced by third generation sequencing, which produces much longer reads without the need for sequence amplification (32), oligo(dT)-primed methods have the distinct advantage of dispensing with the need to sequence the entire RNA molecule, instead focusing solely on mRNA counting to quantify gene expression. In this study, using exclusively previously published publicly accessible datasets, we uncover a heretofore unexplored source of systematic bias in this type of count data, whereby the underlying assumption that the number of cDNA fragments is proportional to the number of mRNA molecules does not hold true for all genes.

Using a dataset with ERCC spike-in control RNA fragments, we show that when non-terminal alignments are removed, the resulting accuracy of the output measurement of known input quantities improves significantly. In the subsequent analysis, we specifically pinpoint internal, genome-encoded A-SNRs as the most likely culprit by showing that there is a significant RNA coverage enrichment upstream of these sequences, and that this enrichment increases with increasing length of these A-SNRs. We then show that in all datasets studied for which a gold standard measurement is available, removal of sequencing read alignments upstream of A-SNRs significantly improves the data accuracy.

Additionally, in all datasets tested, the resulting accuracy after such specific optimized filtering was better than after indiscriminate removal of all non-terminal alignments (i.e., the optimal *minimal A-SNR length* was always higher than zero, see Methods). This is probably the result of the genome reference transcript annotations to date not yet being fully comprehensive. Because of the still incomplete knowledge of alternative polyadenylation sites across the genome, some of the alignments that are labeled non-terminal may in fact be upstream of an unknown transcript 3’ end and their indiscriminate removal (which does not occur during specific removal of alignments upstream of A-SNRs) can lead to sub-optimal data accuracy.

Currently, most oligo(dT)-primed datasets do not have their random oligo-primed counterparts to provide a gold standard measurement. Therefore, in order to correctly filter out internally primed alignments from the data using our algorithm without the gold standard dataset, the optimal values of the filtering parameters (*maximal coverage distance, minimal A-SNR length,* and *maximal number of mismatches*) must be known *a priori*. From our iterative optimization, we found that these parameters differ between datasets, which could be caused by the protocol-specific differences. Outside of this study, these parameters may further vary across different tissue types, research groups, and batches. We therefore propose that obtaining a random oligo-primed measurement from the same sample may be the best way to ensure accuracy of oligo(dT)-primed data.

Alternatively, candidate values of these parameters may be estimated from the results in this study, ensuring that filtering improves the accuracy of the sequencing data, although perhaps not optimally so. In all datasets studied, almost all combinations of the three filtering parameters yielded improvement in data accuracy. One notable exception was the QuantSeq bulk dataset where filtering using values of *minimal A-SNR length* less than nine sometimes yielded decrease in accuracy, whereby decreasing the *maximal number of mismatches* allowed for lower *minimal A-SNR lengths* to be still beneficial to the filtering process. Similarly, in both 10X single cell datasets studied, only combinations of low *maximal coverage distance* and low *minimal A-SNR length* yielded decrease in data accuracy, especially at higher *maximal number of mismatches* allowed. In general, however, our observations suggest that filtering using our algorithm with the *maximal coverage distance* of 300 and the *minimal A-SNR length* of 10 with up to one *mismatch* should be guaranteed to improve the accuracy of any oligo(dT) priming-based dataset.

Future studies may be needed to provide additional clarifications about the nature of this phenomenon and further optimize the data filtration algorithm. For example, in this study, we assumed that internal oligo(dT) priming is most likely to occur on genome-encoded poly(A) sequences with only occasional (up to two) mismatches. While it is well established that thymine nucleotides best hybridize to adenines, their selective preference for the three possible “mismatches” is less clear. For example, a previous study suggests that oligo(dT)s may have a high hybridization rate also onto AG-rich sequences (20). Furthermore, we did not focus on RNA sequencing methods that, following oligo(dT) priming, use full transcript length sequencing to provide gene expression quantification as well as splice variant information, such as SMART-seq. We hypothesize that these methods may be similarly affected by internal priming, although to a different extent.

Nevertheless, the findings in this study clearly demonstrate that caution should be exercised when using oligo(dT) priming-based RNA sequencing library preparation methods. An additional step of data analysis in the form of filtering of the alignments that likely resulted from internal oligo(dT) priming should be carried out especially when directly comparing relative expression of one gene to another or inferring new 3’ untranslated region annotations, with special attention given to the genes we have highlighted as those most likely to be affected by internal priming.

## Methods

### Datasets used

All datasets used originated from previous publications by authors not associated with this study and are publicly available. For the *ERCC spike-in standards analysis*, the QuantSeq 3’ mRNA-Seq REV dataset by Wu, Schmid, Rib et al. (26) was obtained from Gene Expression Omnibus (GEO) Accession GSE137612. Specifically, the sample “siGFP_noPAP_in_batch5” (FASTQ file from Sequence Read Archive (SRA) SRR10134316) was selected due to being a control sample with the highest sequencing depth. This dataset was also used to calculate the *aggregate exonic RNA sequencing coverage* upstream of transcript 3’ ends and A-SNRs, along with the QuantSeq dataset by Ma et al. (5) and PBMC1 10X (v2) A dataset by Ding et al. (28), as described below. *Parameter optimization through iterative filtering* was carried out on the following datasets: QuantSeq 3’ mRNA-Seq FWD dateset by Ma et al. (5) was obtained from GEO Accession GSE116949, using the “Lexogen control diet 1” sample (SRA FASTQ file SRR7510922) along with the associated random oligo priming-based “KAPA control diet 1” sample (SRA FASTQ file SRR7510916). The 10X, CEL-seq2, Drop-Seq, inDrop, and Seq-well single cell (Human PBMC1 and PBMC2) and single nuclei (Mouse Cortex1) datasets by Ding et al. (28) were obtained along with the respective random oligo priming-based bulk datasets from GEO Accession GSE132044.

### ERCC spike-in standards analysis

Using the FASTQ files by Wu, Schmid, Rib et al. (26), a BAM file was generated as described in the original publication except for removal of alignments mapped to genomic A-rich positions. STAR v2.7.3a was used for genome indexing and read alignment (33).

Subsequently, all alignments mapped to the ERCC reference sequences were counted with a custom python script using package pysam v0.16.0.1, while alignments that mapped further than 75 bp upstream of the ERCC poly(A) tails were counted as non-terminal. The distance of 75 bp was selected as a conservative cutoff to signify a distinct priming event, considering that Wu, Schmid, Rib et al. clustered any alignments within 25 bp for the purposes of their 3’ end analysis and the sequencing length used to generate this data was 75 bp. ERCCs with no alignments detected (before removal of non-terminal alignments) were removed from the data. In order to plot and calculate the correlations of input concentrations vs. output alignment counts, the input concentrations (originally in attomoles/ul) were multiplied by 100 to obtain attomoles/100ul. Subsequently, both the input concentrations and the output alignment counts were log-transformed by adding 1 and taking the natural logarithm. Aggregate coverage per bp along the ERCC reference sequences was also obtained using pysam.

### Genome scanning for A-SNRs

We used the GRCh38 v3.0.0 Cell Ranger reference (human), mm10 v3.0.0 Cell Ranger reference (mouse), Dmel_Release_6 FlyBase reference (fruit fly; 34), and Os-Nipponbare-Reference-IRGSP-1.0 reference (rice; 35) for genome scanning and assignment of genomic annotations. The genome was scanned for A-SNRs with a custom python script using Biopython v1.74 Bio.SeqIO package (36) and relative (observed/expected) A-SNR frequency was calculated as described by Murray (27). Genomic annotations were extracted using python package gffutils v0.10.1. The proportions of SNRs at each length assigned to their respective genomic annotations were normalized by the total proportion of the genome with this annotation, as follows: genomic regions annotated as “exon” in at least one transcript were considered exonic, the remaining non-exonic genomic regions covered by at least one transcript (annotated as “transcript” in most references, or as “mRNA” in the fruit fly reference) were considered intronic, and the remaining genomic regions were considered intergenic.

### Aggregate exonic RNA sequencing coverage

Each of the datasets was processed as described by the authors in the respective original publication; datasets by Ding et al. (28) were always processed using their scumi pipeline to ensure fair comparison between different library preparation methods. Each of the respective reference annotations was scanned for transcript 3’ ends, the reference sequences were scanned for A-SNRs, and the dataset BAM files were used to calculate the surrounding RNA sequencing coverage. Per-base sequencing coverage was subsequently normalized by the expected (mean) per-base coverage across all exons (i.e., total exonic coverage across the entire genome divided by the number of bps annotated as exons in at least one expressed transcript) and divided by the number of potential priming events (transcript 3’ ends or A-SNRs).

### Python algorithm to filter out internal oligo(dT)-primed alignments

We designed a custom python algorithm that uses Biopython’s Bio.SeqIO, gffutils, and pysam packages to scan a given reference genome for A-SNRs of given minimal length (the *minimal A-SNR length* parameter) with a given maximal number of mismatches (the *maximal number of mismatches* parameter), in order to remove all non-terminal alignments in a given dataset that are found up to a given coverage distance upstream of each A-SNR identified, but not within the same distance of a 3’ end of any annotated transcript (the *maximal coverage distance* parameter). Specifically, the algorithm does so by carrying out the following sequence of steps:

1. Scan the reference genome annotation (a GTF/GFF file) for all transcripts and save those that have any exonic coverage associated with them in the sequencing dataset (a BAM file).
2. Protect (i.e., remove) the 3’-terminal segments of all covered transcripts up to exonic (i.e., not counting introns) length equal to the *maximal coverage distance*. Any sequencing reads aligned within these protected segments will be kept in the dataset.
3. Scan the reference genome sequence (a FASTA file) to identify all A-SNRs whose length is equal to or greater than the *minimal A-SNR length*, with the *maximal number of mismatches* allowed. No mismatches are allowed in the first or last bp of the A-SNR.
4. Identify internal priming segments up to the *maximal coverage distance* upstream of these A-SNRs and overlapping non-terminal exonic portions of all covered transcripts.
5. Remove all alignments in the dataset (a BAM file) that overlap with these internal priming segments.

It is also possible to run this algorithm with the option to include the intronic annotations and coverage (as was done in the case of single nuclei RNA sequencing data filtering). When *minimal A-SNR length* is set to 0 (or “None”), this algorithm removes all non-terminal alignments – i.e., all reads that are not aligned within the segments protected in step 2., regardless of their association with A-SNRs.

### Parameter optimization through iterative filtering

Each of the oligo(dT) priming-based datasets studied was processed as described in the respective original publication except for additionally filtering thus generated BAM files, as described below, before further processing the data.

Using the algorithm described in the previous section and varying the three input parameters, we repeatedly filtered each of the oligo(dT) priming-based datasets studied to maximize the correlation with the random oligo priming-based gold standard dataset obtained from the same biological sample (the correlations were calculated as described in the respective original publications). Due to the limitations of the computational resources available, instead of testing all the possible input parameter combinations, we carried out the filtering iteratively, using the set of parameters in the vicinity of those that previously yielded the highest correlation, assuming that the underlying function was generaly concave with one global maximum and no local maxima.

We selected a conservative set of starting filtering parameters based on previously used internal priming filtration criteria (see Introduction) as well as our own results (Figure 2CD), such that the optimal values were expected to be found within the ranges tested, as follows:

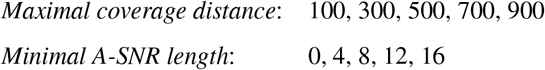

The *minimal A-SNR length* of “0” indicates that all nonterminal alignments were removed, regardless of their association with any A-SNRs. Filtering using all 25 possible combinations of these starting parameter values yielded a 5 × 5 matrix of resulting correlations (Figure S3). The parameter values yielding the highest correlation were then selected to be the central points in the subsequently tested value ranges of these parameters. For example, among the parameters used above, if the highest correlation was found by using the *maximal coverage distance* of 300 and *minimal A-SNR length* of 4, the subsequently tested values were:

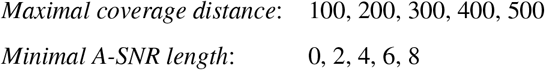

This process was repeated until maximal resolution was achieved and no further iterations were possible. Such optimization was carried out for sequential values of *maximal number of mismatches* starting with zero, until no further improvement in correlation was observed.

### Ranking genes by their likelihood to exhibit internal oligo(dT) priming

Optimally filtered 10X PBMC1 dataset by Ding et al. (28) was used to rank the genes. We used R package zoib (37) to derive the per-gene dependence of proportion of optimally filtered out alignments on the genes’ A-SNR content and length, using a Bayesian zero-one inflated beta regression model, as follows:

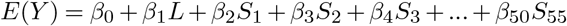

where *E*(*Y*) ∈ [0,1] is the expected proportion of internally primed reads removed by filtering, *L* is the length of the gene in bp, and *S_N_* is the number of exonic A-SNRs of length *N* (including up to 1 mismatch) contained in the given gene. Note that not all A-SNR lengths below 55 were represented in the model due to their absence among exonic A-SNRs across the expressed genes used to derive this model. The specific model used was selected as the one that generated the lowest mean squared error (MSE) on out-of-sample data among all tested models. All human genes were subsequently rank-ordered by the resulting *E*(*Y*) prediction.

## Code availability

The python code for the BAM filtering algorithm as well as all other analyses described in this publication is available at https://github.com/MarekSvob/polyAfilter.

## ACKNOWLEDGEMENTS

This research was supported by the Rosaline Borison Memorial Fund and the Burroughs Wellcome Fund. We would like to thank Ding et al. (28), especially J. Ding and S. K. Simmons, for their responsiveness during the communication of the details and clarifications that enabled replication of some of their data analyses. This article was written using the HenriquesLab bioRxiv template.

## Supplementary Material

**Figure S1.**
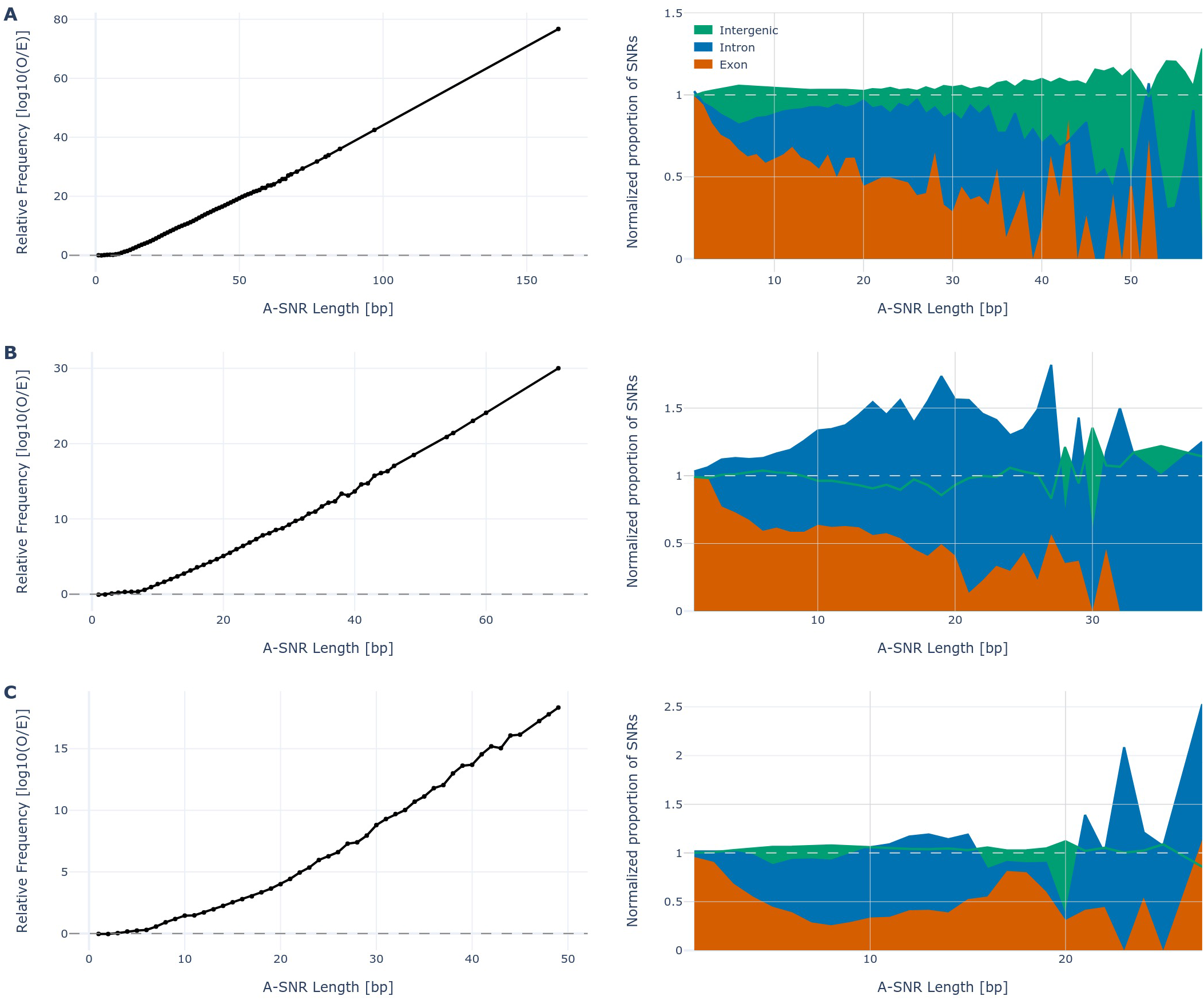
The relative frequencies of A-SNRs across various genome references. A-SNR analysis for reference genomes of *Mus musculus* (**A.**), *Drosophila melanogaster* (**B.**), and *Oriza sativa* (**C.**). Left: Log10 of relative (observed/expected) frequency of A-SNRs by length across the entire respective reference. The expected frequency was calculated based on the probability that A-SNRs of given length be found by pure chance. Right: The relative proportion of A-SNRs of each length by their genomic annotations, normalized by the respective proportions of the genome thus annotated. SNR lengths represented by fewer than 10 SNRs were removed. Color legend from **A.** applies to **B.** and **C.** as well. Gray dashed line in each plot represents the expected value.

**Figure S2.**
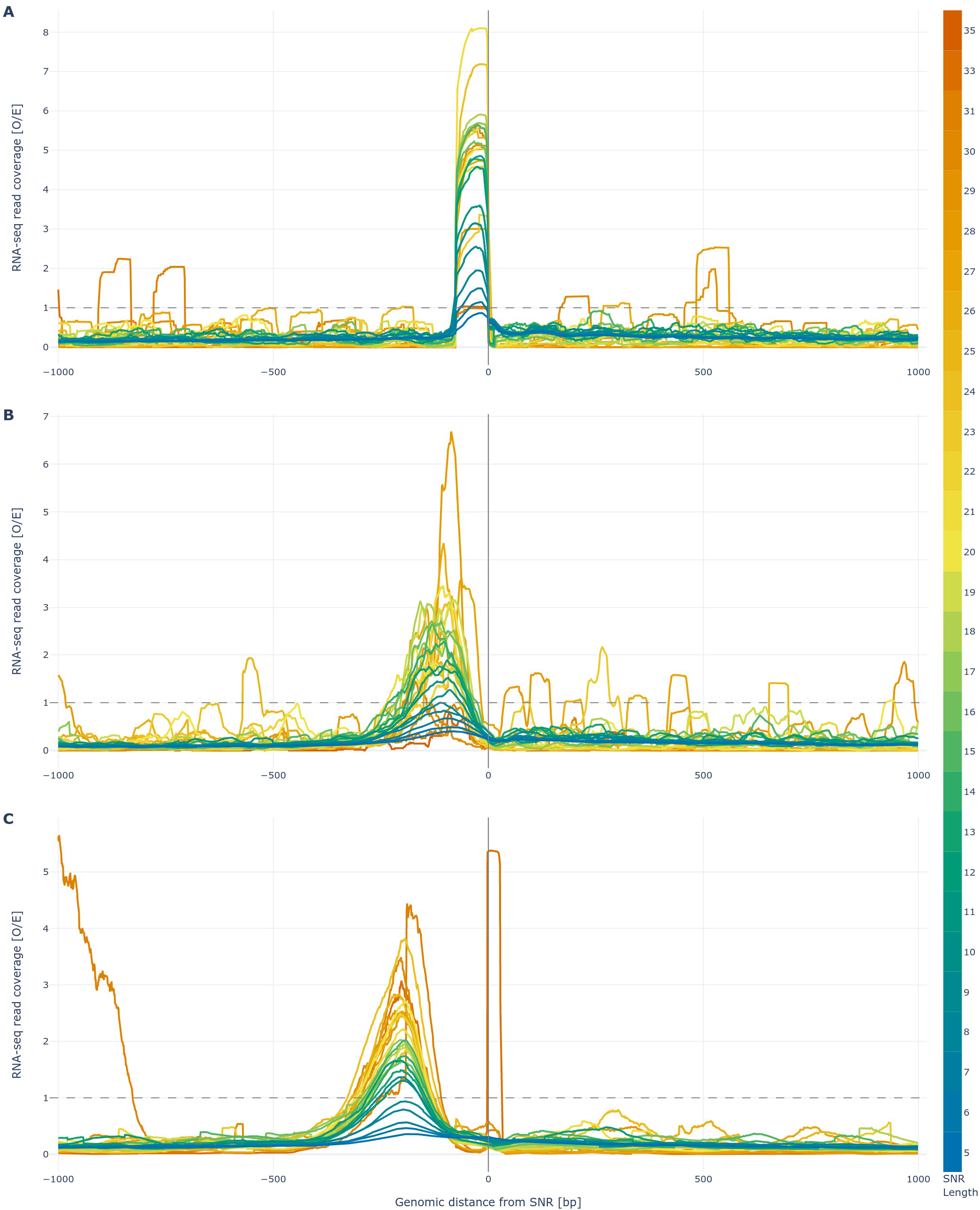
Sequencing coverage in the vicinity of SNRs by length. Normalized aggregate stranded exonic RNA sequencing coverage in the vicinity of A-SNRs of five nucleotides or longer, grouped by A-SNR length depicted by the color scale. The starts (5’ ends) of all A-SNRs are aligned at “0 bp” in the sense orientation. Sequencing coverage from datasets by Wu, Schmid, Rib et al. (“Lexo REV”, **A.**; 26), Ma et al. (“Lexo FWD”, **B.**; 5), and Ding et al. (“10X”, **C.**; 28) is depicted. In each graph, the gray dashed line represents the expected value if exonic coverage was randomly (evenly) distributed along exons. A-SNR outliers with associated coverage higher than 100 × the expected value and A-SNR lengths represented by fewer than 10 A-SNRs were removed from this data.

**Figure S3.**
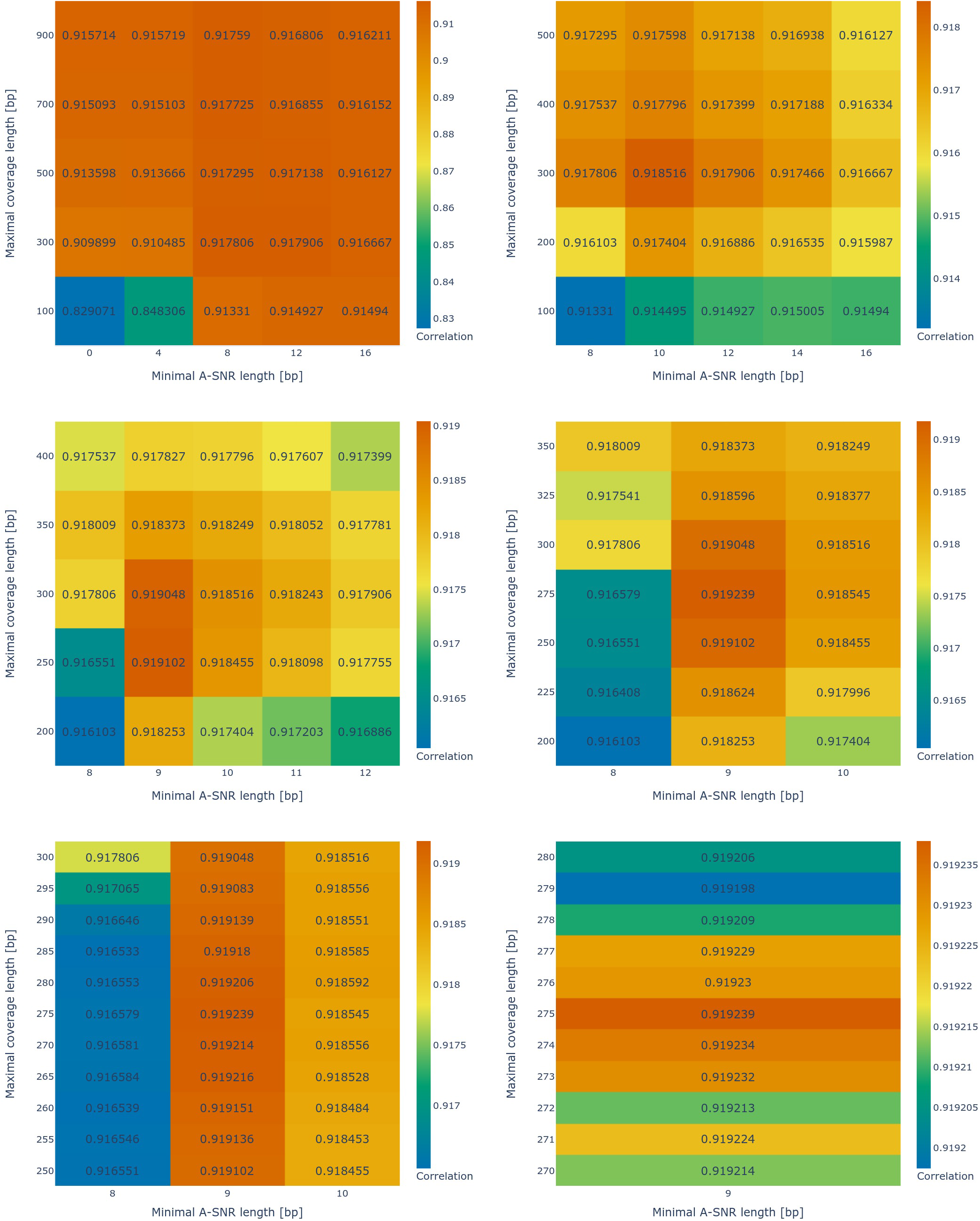
An example of parameter optimization through stepwise alignment filtering. Sequential steps in filtering optimization of the PBMC1 10X (v2) dataset by Ding et al. (28), at varying values of *minimal A-SNR length* and *maximal coverage distance*, with the *maximal number of mismatches* equal to one. Each subsequent set of parameters centers around those with the highest correlation(s) obtained at the previous step. Correlations with the associated bulk dataset in each heatmap are indicated by the number and color from the adjacent color scale.

**Figure S4.**
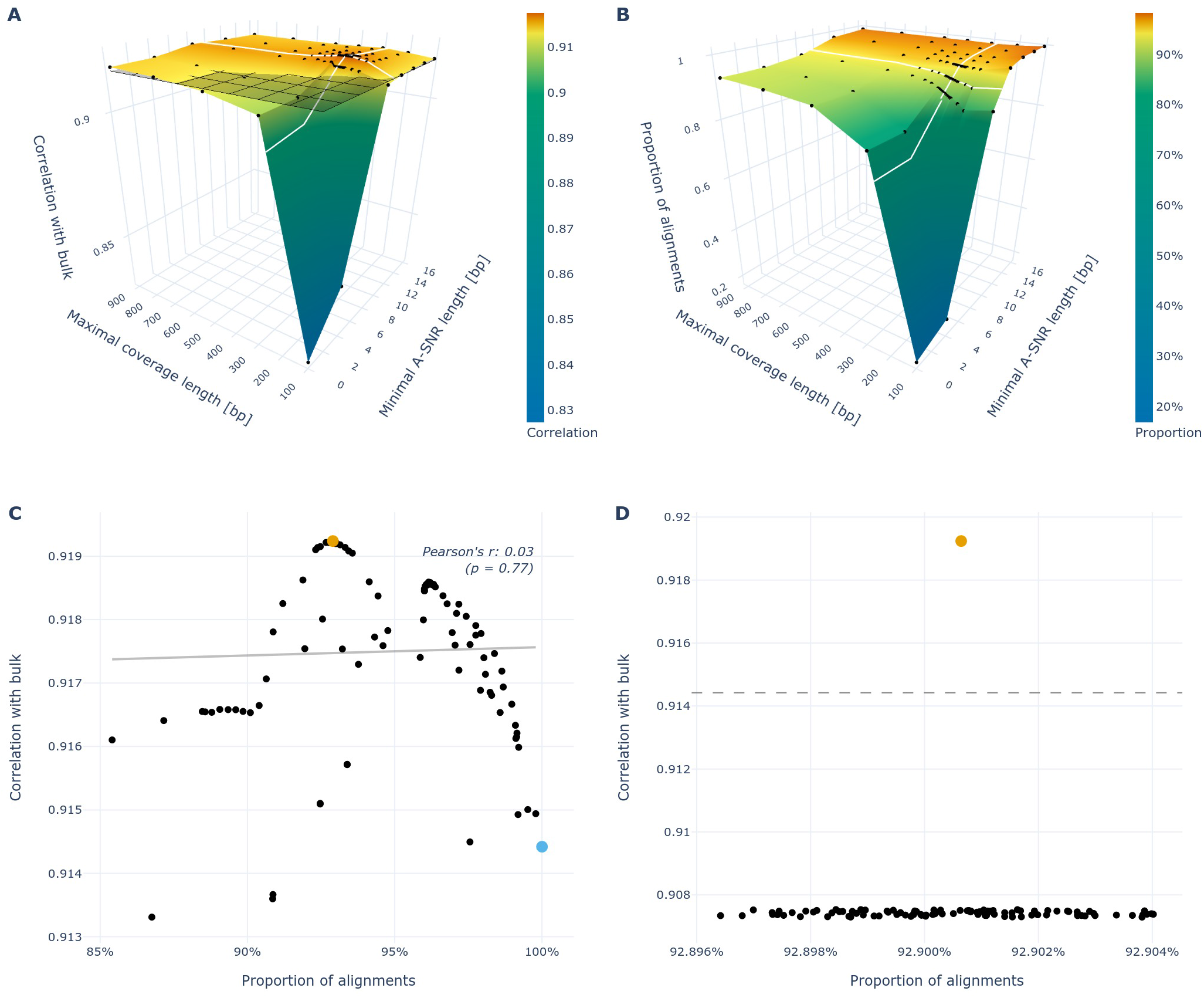
The correlation with bulk vs. the proportion of exonic alignments remaining after filtering. **A.** A 3D surface representation of all the correlations from Figure S3. The dark gray opaque plane with black orthogonal lines indicates the correlation of the original PBMC1 10X (v2) dataset by Ding et al. (28) with the associated bulk dataset before filtering. **B.** A 3D surface representation of the remaining exonic alignment proportions after filtering for each respective combination of *minimal A-SNR length* and *maximal coverage distance* values from **A.**, with the value before filtering being equal to one (not shown). In both **A.** and **B.**, the measured values are indicated as black dots with the surface interpolated in between and the white lines indicate the optimal *minimal A-SNR length* and *maximal coverage distance* whose combination yielded the highest correlation value after filtering. **C.** A plot of correlations from **A.** vs. proportions from **B.** shows the lack of correlation (line of best fit in solid gray) between the two. Four outliers with proportions of exonic reads lower than 0.85 after filtering were removed. The blue dot represents the original dataset before filtering. **D.** A plot of the remaining exonic alignment proportions vs. resulting correlations after 100 times randomly filtering out the same proportion of exonic alignments as in the optimally filtered dataset (~7.1%; note that the slight variations in the final proportions are caused by varying the seed to initiate the pseudorandom filtering). The dashed gray line indicates the correlation before filtering. In both **C.** and **D.**, the optimized dataset is represented by the orange dot.

**Table S1.** The numbers of intragenic alignments in each dataset’s BAM file before and after optimal filtering. The percentages indicate the proportion of the respective alignment types remaining after filtering.

| Method | Sample | Introns Included | Before Filtering |  | After Filtering |  |
| --- | --- | --- | --- | --- | --- | --- |
|  |  |  | Exonic | Intronic | Exonic | Intronic |
| QuantSeq | Mouse liver (bulk) | No | 14 058 019 | 1 562 763 | 13 527 504<br>(96.2%) | 1 562 763<br>(100.0%) |
| 10X | Human PBMC1 (sc) | No | 220 587 386 | 102 312 104 | 204 927 102<br>(92.9%) | 102 312 104<br>(100.0%) |
| 10X | Human PBMC2 (sc) | No | 176 044 691 | 93 636 640 | 156 127 308<br>(88.7%) | 93 636 640<br>(100.0%) |
| CEL-Seq2 | Human PBMC1 (sc) | No | 173 405 676 | 432 682 735 | 72 491 147<br>(41.8%) | 432 682 735<br>(100.0%) |
| Drop-Seq | Human PBMC1 (sc) | No | 155 733 230 | 62 564 219 | 70 809 314<br>(45.5%) | 62 564 219<br>(100.0%) |
| inDrop | Human PBMC1 (sc) | No | 165 292 779 | 104 599 750 | 131 549 685<br>(79.6%) | 104 599 750<br>(100.0%) |
| Seq-Well | Human PBMC1 (sc) | No | 70 258 471 | 48 897 396 | 24 399 738<br>(34.7%) | 48 897 396<br>(100.0%) |
| 10X | Mouse Cortex1 (sn) | Yes | 121 416 773 | 133 732 777 | 89 931 124<br>(74.1%) | 696 713<br>(0.5%) |
| 10X | Mouse Cortex1 (sn) | No | 121 416 773 | 133 732 777 | 93 787 604<br>(77.2%) | 133 732 777<br>(100.0%) |

**Figure S5.**
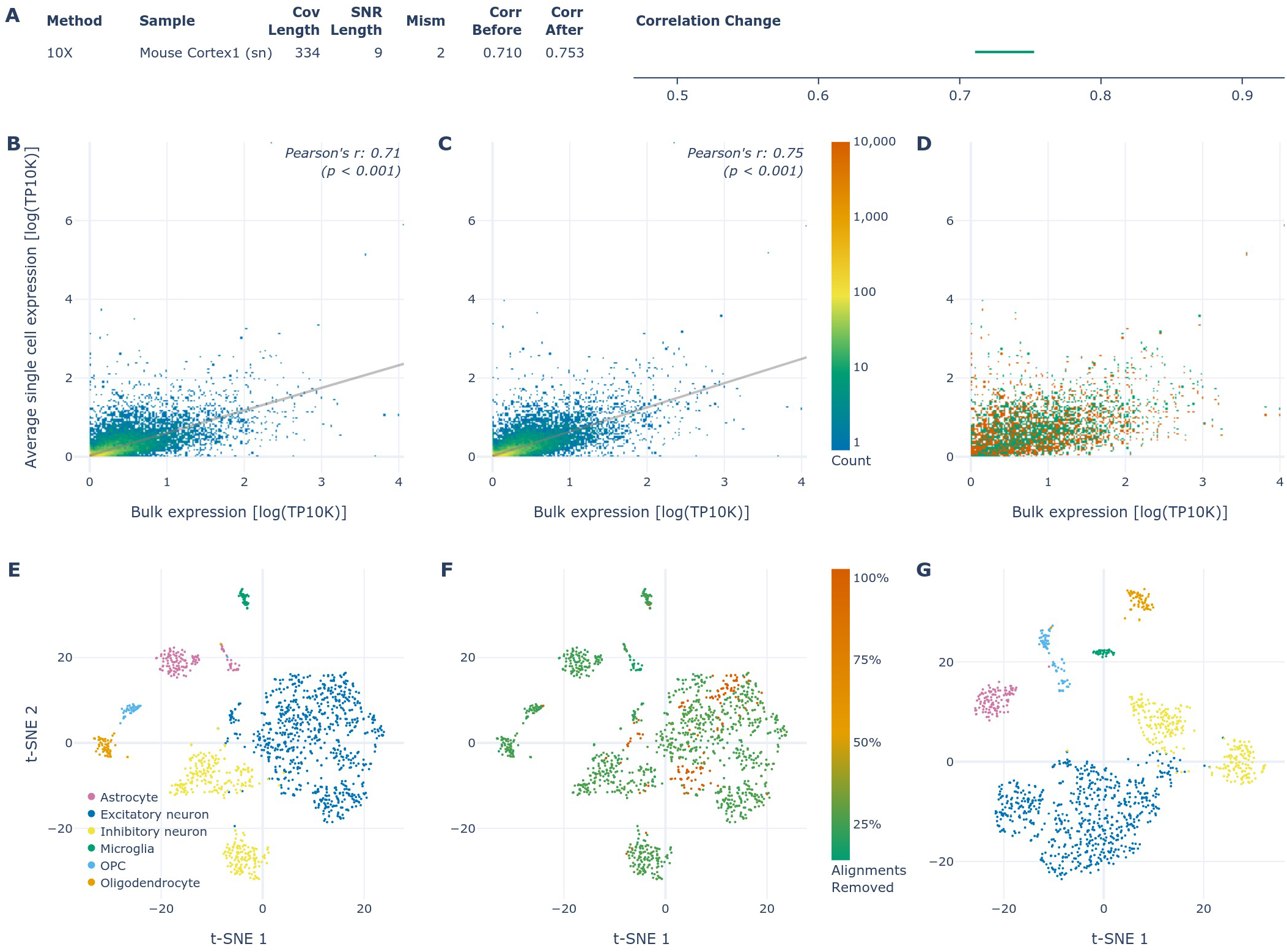
Optimized filtering of the single nuclei dataset with only exonic alignments included in gene expression quantification. **A.** Table of the optimized filtering values for the Mouse Cortex1 single nuclei 10X dataset by Ding et al. (28) when intronic alignments are excluded from gene expression quantification (in both the single nuclei and the associated bulk datasets). Correlation change between before and after optimal filtering visualized on the same x-axis as in Figure 3); *p* << 0.001 for each correlation, as well as for the difference between correlations before and after filtering, adjusted using the Bonferroni correction. Abbreviations used: “Cov Length”: maximal coverage distance; “SNR Length”: minimal A-SNR length; “Mism”: maximal number of mismatches; “Corr”: correlation; “sn”: single nuclei. **B.** Correlation of gene expression between the 10X (oligo(dT)) and bulk (random oligos) methods carried out on the same sample (mouse cortex single nuclei). **C.** Correlation between the same datasets as in **B.** after filtering the oligo(dT) dataset using the optimal parameters listed in **A.** Both **B.** and **C.** are 2D histograms sharing the rainbow color scale depicting the density of genes in each region with the line of best fit in gray. **D.** Visualization of changes between **B.** and **C.**, where the regions with fewer and more genes in **C.** relative to **B.** are depicted by orange and green, respectively. **E.** t-SNE plot of the single nuclei mouse cortex 10X dataset colored by the cell types detected. **F.** The same t-SNE plot as in **E.**, colored by the proportion of alignments optimally filtered out from each cell, as per the adjacent color scale. **G.** t-SNE plot of the same dataset as in **E.** and **F.**, after the internally primed alignments have been optimally filtered out. The colors correspond to the same cell types as in **E.**

**Table S2.**
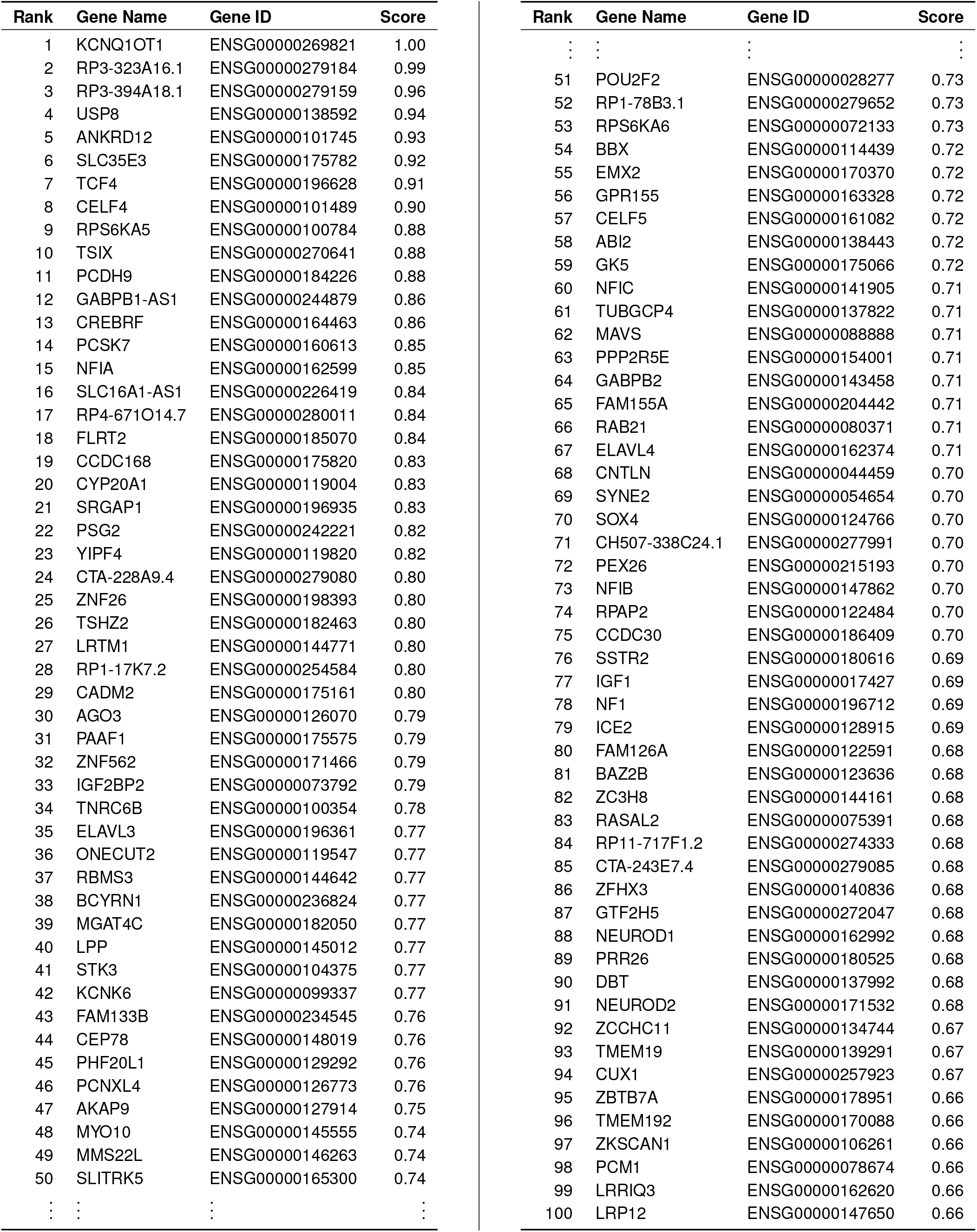
The top 100 human genes whose poly(dT)-based RNA sequencing expression quantification is predicted to be most likely affected by internal priming. The score represents the proportion of exonic alignments predicted by the linear model to originate from internal priming.

## Notes

https://github.com/MarekSvob/polyAfilter

https://www.ncbi.nlm.nih.gov/geo/query/acc.cgi?acc=GSE137612

https://www.ncbi.nlm.nih.gov/geo/query/acc.cgi?acc=GSE116949

https://www.ncbi.nlm.nih.gov/geo/query/acc.cgi?acc=GSE132044

### Competing Interest Statement

The authors have declared no competing interest.

## Bibliography

1. Alexander F. Palazzo and Eliza S. Lee. Non-coding RNA: What is functional and what is junk? Frontiers in Genetics, 6:2, jan 2015. ISSN 16648021. doi:10.3389/fgene.2015.00002.

2. Peng Cui, Qiang Lin, Feng Ding, Chengqi Xin, Wei Gong, Lingfang Zhang, Jianing Geng, Bing Zhang, Xiaomin Yu, Jin Yang, Songnian Hu, and Jun Yu. A comparison between ribo-minus RNA-sequencing and polyA-selected RNA-sequencing. Genomics, 96(5):259–265, nov 2010. ISSN 08887543. doi:10.1016/j.ygeno.2010.07.010.

3. Shanrong Zhao, Ying Zhang, Ramya Gamini, Baohong Zhang, and David Von Schack. Evaluation of two main RNA-seq approaches for gene quantification in clinical RNA sequencing: PolyA+ selection versus rRNA depletion. Scientific Reports, 8(1):4781, dec 2018. ISSN 20452322. doi:10.1038/s41598-018-23226-4.

4. Savita Shanker, Ariel Paulson, Howard J. Edenberg, Allison Peak, Anoja Perera, Yuriy O. Alekseyev, Nicholas Beckloff, Nathan J. Bivens, Robert Donnelly, Allison F. Gillaspy, Deborah Grove, Weikuan Gu, Nadereh Jafari, Joanna S. Kerley-Hamilton, Robert H. Lyons, Clifford Tepper, and Charles M. Nicolet. Evaluation of commercially available RNA amplification kits for RNA sequencing using very low input amounts of total RNA. Journal of Biomolecular Techniques, 26(1):4–18, 2015. ISSN 19434731. doi:10.7171/jbt.15-2601-001.

5. Feiyang Ma, Brie K. Fuqua, Yehudit Hasin, Clara Yukhtman, Chris D. Vulpe, Aldons J. Lusis, and Matteo Pellegrini. A comparison between whole transcript and 3’ RNA sequencing methods using Kapa and Lexogen library preparation methods. BMC Genomics, 20(1):9, jan 2019. ISSN 14712164. doi:10.1186/s12864-018-5393-3.

6. J Eberwine, H Yeh, K Miyashiro, Y Cao, S Nair, R Finnell, M Zettel, and P Coleman. Analysis of gene expression in single live neurons. Proceedings of the National Academy of Sciences of the United States of America, 89(7):3010–4, apr 1992. ISSN 0027-8424. doi:10.1073/PNAS.89.7.3010.

7. Evan Z Macosko, Anindita Basu, Rahul Satija, James Nemesh, Karthik Shekhar, Melissa Goldman, Itay Tirosh, Allison R Bialas, Nolan Kamitaki, Emily M Martersteck, John J Trombetta, David A Weitz, Joshua R Sanes, Alex K Shalek, Aviv Regev, and Steven A McCarroll. Highly Parallel Genomewide Expression Profiling of Individual Cells Using Nanoliter Droplets. Cell, 161 (5):1202–1214, may 2015. ISSN 1097-4172. doi:10.1016/j.cell.2015.05.002.

8. Allon M. Klein, Linas Mazutis, Ilke Akartuna, Naren Tallapragada, Adrian Veres, Victor Li, Leonid Peshkin, David A. Weitz, and Marc W. Kirschner. Droplet Barcoding for Single-Cell Transcriptomics Applied to Embryonic Stem Cells. Cell, 161(5):1187–1201, may 2015. ISSN 0092-8674. doi:10.1016/J.CELL.2015.04.044.

9. Grace X.Y. Zheng, Jessica M. Terry, Phillip Belgrader, Paul Ryvkin, Zachary W. Bent, Ryan Wilson, Solongo B. Ziraldo, Tobias D. Wheeler, Geoff P. McDermott, Junjie Zhu, Mark T. Gregory, Joe Shuga, Luz Montesclaros, Jason G. Underwood, Donald A. Masquelier, Stefanie Y. Nishimura, Michael Schnall-Levin, Paul W. Wyatt, Christopher M. Hindson, Rajiv Bharadwaj, Alexander Wong, Kevin D. Ness, Lan W. Beppu, H. Joachim Deeg, Christopher McFarland, Keith R. Loeb, William J. Valente, Nolan G. Ericson, Emily A. Stevens, Jerald P. Radich, Tarjei S. Mikkelsen, Benjamin J. Hindson, and Jason H. Bielas. Massively parallel digital transcriptional profiling of single cells. Nature Communications, 8:14049, jan 2017. ISSN 20411723. doi:10.1038/ncomms14049.

10. Todd M Gierahn, Marc H Wadsworth, Travis K Hughes, Bryan D Bryson, Andrew Butler, Rahul Satija, Sarah Fortune, J Christopher Love, and Alex K Shalek. Seq-Well: portable, low-cost RNA sequencing of single cells at high throughput. Nature Methods, 14:395–398, feb 2017. ISSN 1548-7105. doi:10.1038/nmeth.4179.

11. Junyue Cao, Jonathan S Packer, Vijay Ramani, Darren A Cusanovich, Chau Huynh, Riza Daza, Xiaojie Qiu, Choli Lee, Scott N Furlan, Frank J Steemers, Andrew Adey, Robert H Waterston, Cole Trapnell, and Jay Shendure. Comprehensive single-cell transcriptional profiling of a multicellular organism. Science (New York, N.Y.), 357(6352):661–667, aug 2017. ISSN 1095-9203. doi:10.1126/science.aam8940.

12. Michael Hagemann-Jensen, Christoph Ziegenhain, Ping Chen, Daniel Ramsköld, Gert Jan Hendriks, Anton J.M. Larsson, Omid R. Faridani, and Rickard Sandberg. Single-cell RNA counting at allele and isoform resolution using Smart-seq3. Nature Biotechnology, 38:708–714, jun 2020. ISSN 15461696. doi:10.1038/s41587-020-0497-0.

13. Tamar Hashimshony, Naftalie Senderovich, Gal Avital, Agnes Klochendler, Yaron de Leeuw, Leon Anavy, Dave Gennert, Shuqiang Li, Kenneth J Livak, Orit Rozenblatt-Rosen, Yuval Dor, Aviv Regev, and Itai Yanai. CEL-Seq2: sensitive highly-multiplexed single-cell RNA-Seq. Genome Biology, 17(77), 2016. doi:10.1186/s13059-016-0938-8.

14. Pamela Moll, Michael Ante, Alexander Seitz, and Torsten Reda. QuantSeq 3’ mRNA sequencing for RNA quantification. Nature Methods, 11([advertising feature]):i–iii, 2014. ISSN 1548-7091. doi:10.1038/nmeth.f.376.

15. Brian K. Lohman, Jesse N. Weber, and Daniel I. Bolnick. Evaluation of TagSeq, a reliable low-cost alternative for RNAseq. Molecular Ecology Resources, 16 (6):1315–1321, nov 2016. ISSN 1755-0998. doi:10.1111/1755-0998.12529.

16. Gabriel Sholder, Thomas A. Lanz, Robert Moccia, Jie Quan, Estel Aparicio-Prat, Robert Stanton, and Hualin S. Xi. 3’Pool-seq: an optimized cost-efficient and scalable method of whole-transcriptome gene expression profiling. BMC Genomics, 21(64), jan 2020. ISSN 1471-2164. doi:10.1186/S12864-020-6478-3.

17. Douglas Kyung Nam, Sanggyu Lee, Guolin Zhou, Xiaohong Cao, Clarence Wang, Terry Clark, Jianjun Chen, Janet D. Rowley, and San Ming Wang. Oligo(dT) primer generates a high frequency of truncated cDNAs through internal poly(A) priming during reverse transcription. Proceedings of the National Academy of Sciences of the United States of America, 99(9):6152–6156, apr 2002. ISSN 00278424. doi:10.1073/pnas.092140899.

18. Haibo Zhang, Jun Hu, Michael Recce, and Bin Tian. PolyA_DB: A database for mammalian mRNA polyadenylation. Nucleic Acids Research, 33(Issue suppl_1):D116–D120, 2005. ISSN 03051048. doi:10.1093/nar/gki055.

19. Ju Youn Lee, Ijen Yeh, Ji Yeon Park, and Bin Tian. PolyA_DB 2: mRNA polyadenylation sites in vertebrate genes. Nucleic Acids Research, 35(Issue suppl_1):D165–D168, 2007. ISSN 03051048. doi:10.1093/nar/gkl870.

20. Joel H. Graber, Fathima I. Nazeer, Pei Chun Yeh, Jason N. Kuehner, Sneha Borikar, Derick Hoskinson, and Claire L. Moore. DNA damage induces targeted, genome-wide variation of poly(A) sites in budding yeast. Genome Research, 23(10):1690–1703, oct 2013. ISSN 10889051. doi:10.1101/gr.144964.112.

21. Stefan Wilkening, Vicent Pelechano, Aino I. Järvelin, Manu M. Tekkedil, Simon Anders, Vladimir Benes, and Lars M. Steinmetz. An efficient method for genome-wide polyadenylation site mapping and RNA quantification. Nucleic Acids Research, 41(5):e65, mar 2013. ISSN 13624962. doi:10.1093/nar/gks1249.

22. Kevin Roy, Jason Gabunilas, Abigail Gillespie, Duy Ngo, and Guillaume F. Chanfreau. Common genomic elements promote transcriptional and DNA replication roadblocks. Genome Research, 26(10):1363–1375, oct 2016. ISSN 15495469. doi:10.1101/gr.204776.116.

23. Gioele La Manno, Ruslan Soldatov, Amit Zeisel, Emelie Braun, Hannah Hochgerner, Viktor Petukhov, Katja Lidschreiber, Maria E. Kastriti, Peter Lön-nerberg, Alessandro Furlan, Jean Fan, Lars E. Borm, Zehua Liu, David van Bruggen, Jimin Guo, Xiaoling He, Roger Barker, Erik Sundström, Gonçalo Castelo-Branco, Patrick Cramer, Igor Adameyko, Sten Linnarsson, and Peter V. Kharchenko. RNA velocity of single cells. Nature, 560:494–498, aug 2018. ISSN 14764687. doi:10.1038/s41586-018-0414-6.

24. Oriya Vardi, Inbal Shamir, Elisheva Javasky, Alon Goren, and Itamar Simon. Biases in the SMART-DNA library preparation method associated with genomic poly dA/dT sequences. PLOS ONE, 12(2):e0172769, feb 2017. ISSN 1932-6203. doi:10.1371/JOURNAL.PONE.0172769.

25. Lichun Jiang, Felix Schlesinger, Carrie A. Davis, Yu Zhang, Renhua Li, Marc Salit, Thomas R. Gingeras, and Brian Oliver. Synthetic spike-in standards for RNA-seq experiments. Genome Research, 21(9):1543–1551, sep 2011. ISSN 10889051. doi:10.1101/gr.121095.111.

26. Guifen Wu, Manfred Schmid, Leonor Rib, Patrik Polak, Nicola Meola, Albin Sandelin, and Torben Heick Jensen. A Two-Layered Targeting Mechanism Underlies Nuclear RNA Sorting by the Human Exosome. Cell Reports, 30(7): 2387–2401.e5, feb 2020. ISSN 22111247. doi:10.1016/j.celrep.2020.01.068.

27. Vincent Murray. The frequency of poly(G) tracts in the human genome and their use as a sensor of DNA damage. Computational Biology and Chemistry, 54:13–17, 2015. ISSN 14769271. doi:10.1016/j.compbiolchem.2014.11.006.

28. Jiarui Ding, Xian Adiconis, Sean K. Simmons, Monika S. Kowalczyk, Cynthia C. Hession, Nemanja D. Marjanovic, Travis K. Hughes, Marc H. Wadsworth, Tyler Burks, Lan T. Nguyen, John Y.H. Kwon, Boaz Barak, William Ge, Amanda J. Kedaigle, Shaina Carroll, Shuqiang Li, Nir Hacohen, Orit Rozenblatt-Rosen, Alex K. Shalek, Alexandra Chloé Villani, Aviv Regev, and Joshua Z. Levin. Systematic comparison of single-cell and single-nucleus RNA-sequencing methods. Nature Biotechnology, 38(6):737–746, jun 2020. ISSN 15461696. doi:10.1038/s41587-020-0465-8.

29. Lexogen. QuantSeq 3’ mRNA-Seq Library Prep Kit FWD for Illumina, 2020. https://www.lexogen.com/quantseq-3mrna-sequencing/ (Accessed: 2021-07-17).

30. Trygve E. Bakken, Rebecca D. Hodge, Jeremy A. Miller, Zizhen Yao, Thuc Nghi Nguyen, Brian Aevermann, Eliza Barkan, Darren Bertagnolli, Tamara Casper, Nick Dee, Emma Garren, Jeff Goldy, Lucas T. Graybuck, Matthew Kroll, Roger S. Lasken, Kanan Lathia, Sheana Parry, Christine Rimorin, Richard H. Scheuermann, Nicholas J. Schork, Soraya I. Shehata, Michael Tieu, John W. Phillips, Amy Bernard, Kimberly A. Smith, Hongkui Zeng, Ed S. Lein, and Bosiljka Tasic. Single-nucleus and single-cell transcriptomes compared in matched cortical cell types. PLOS ONE, 13(12):e0209648, dec 2018. ISSN 1932-6203. doi:10.1371/journal.pone.0209648.

31. Jean-Charles Boisset, Judith Vivié, Dominic Grün, Mauro J. Muraro, Anna Lyubimova, and Alexander van Oudenaarden. Mapping the physical network of cellular interactions. Nature Methods, 15(7):547–553, jul 2018. ISSN 1548-7091. doi:10.1038/s41592-018-0009-z.

32. Huanle Liu, Oguzhan Begik, Morghan C. Lucas, Jose Miguel Ramirez, Christopher E. Mason, David Wiener, Schraga Schwartz, John S. Mattick, Martin A. Smith, and Eva Maria Novoa. Accurate detection of m6A RNA modifications in native RNA sequences. Nature Communications 2019 10:1, 10(1):1–9, sep 2019. ISSN 2041-1723. doi:10.1038/s41467-019-11713-9.

33. Alexander Dobin, Carrie A. Davis, Felix Schlesinger, Jorg Drenkow, Chris Zaleski, Sonali Jha, Philippe Batut, Mark Chaisson, and Thomas R. Gingeras. STAR: ultrafast universal RNA-seq aligner. Bioinformatics, 29(1):15–21, jan 2013. ISSN 1367-4803. doi:10.1093/BIOINFORMATICS/BTS635.

34. Gilberto dos Santos, Andrew J. Schroeder, Joshua L. Goodman, Victor B. Strelets, Madeline A. Crosby, Jim Thurmond, David B. Emmert, William M. Gelbart, and the FlyBase Consortium. FlyBase: introduction of the Drosophila melanogaster Release 6 reference genome assembly and large-scale migration of genome annotations. Nucleic Acids Research, 43(D1):D690–D697, jan 2015. ISSN 0305-1048. doi:10.1093/NAR/GKU1099.

35. Nancy A. Eckardt. Sequencing the Rice Genome. The Plant Cell, 12(11): 2011–2017, nov 2000. ISSN 1040-4651. doi:10.1105/TPC.12.11.2011.

36. Peter J. A. Cock, Tiago Antao, Jeffrey T. Chang, Brad A. Chapman, Cymon J. Cox, Andrew Dalke, Iddo Friedberg, Thomas Hamelryck, Frank Kauff, Bartek Wilczynski, and Michiel J. L. de Hoon. Biopython: freely available Python tools for computational molecular biology and bioinformatics. Bioinformatics, 25(11):1422–1423, jun 2009. ISSN 1367-4803. doi:10.1093/BIOINFORMATICS/BTP163.

37. Fang Liu and Yunchuan Kong. zoib: An R package for Bayesian inference for beta regression and Zero/one inflated beta regression. R Journal, 7(2):34–51, 2015. ISSN 20734859. doi:10.32614/rj-2015-019.

